# Mechanical Mapping of Bioprinted Hydrogel Models by Brillouin Microscopy

**DOI:** 10.1101/2021.02.18.431535

**Authors:** Hadi Mahmodi, Alberto Piloni, Robert Utama, Irina Kabakova

## Abstract

Three-dimensional (3D) bioprinting has revolutionised the field of biofabrication by delivering precise, cost-effective and a relatively simple way of engineering *in vitro* living systems in high volume for use in tissue regeneration, biological modelling, drug testing and cell-based diagnostics. The complexity of modern bioprinted systems requires quality control assessment to ensure the resulting product meets the desired criteria of structural design, micromechanical performance and long-term durability. Brillouin microscopy could be an excellent solution for micromechanical assessment of the bioprinted models during or post-fabrication since this technology is non-destructive, label-free and is capable of microscale 3D imaging. In this work, we demonstrate the application of Brillouin microscopy to 3D imaging of hydrogel microstructures created through drop-on-demand bioprinting. In addition, we show that this technology can resolve variations between mechanical properties of the gels with slightly different polymer fractions. This work confirms that Brillouin microscopy can be seen as a characterisation technology complementary to bioprinting, and in the future can be combined within the printer design to achieve simultaneous real-time fabrication and micromechanical characterisation of *in vitro* biological systems.

## 1. Introduction

The emergence of three-dimensional (3D) bioprinting has greatly stimulated the vibrant field of biofabrication by creating new opportunities for engineering *in vitro* living systems for use in tissue regeneration, biological modelling, and cell-based diagnostics [1-3]. 3D bioprinting is an additive manufacturing technique that is based on the deposition of cell-laden biomaterials to form artificial micro-to macroscale structures comparable to living organs and tissues [4, 5]. The most common modalities for tissue fabrication are extrusion and drop-on-demand bioprinting, in which living constructs are additively manufactured via layer-by-layer or, alternatively, droplet-by-droplet deposition of bio-inks [4]. Thanks to controlled and precise deposition, cost-effectiveness and relative simplicity of these technologies, the number of 3D bioprinting applications have been increasing constantly over the past few years. This rapid technological development has recently culminated in stellar demonstrations of the first integrated organs, such as thyroid gland [6] and an ovary [7].

The functionality of living tissues is governed predominantly by their complex architecture [8]. The topographical and mechanical cues provided by the extracellular matrix, together with the specific distribution of morphogens and biochemical signals, are well recognised as major determinants of cell fate both *in vitro* and *in vivo* [9, 10]. With increasing complexity of 3D bioprinting models and diversity in bio-ink materials and printing technology, quality control and testing of bioprinted models becomes the crucial component of biofabrication. Special consideration must be paid to mechanical properties of bioprinted constructs, their long-term stability and integrity. Due to the three-dimensional nature of bioprinted tissue models, which are typically immersed in medium to avoid dehydration, it is often challenging to fully characterise architectural and mechanical properties of these models in a non-destructive manner. For example, shear stress and strain properties of biomaterials can be assessed pre-fabrication by standard micro-rheology [11-13]. These findings, however, have limited significance post-fabrication, since the bioprinting process itself modifies material physical properties by additional pressure, light and/or temperature manipulation. Molecular exchange between matrix and cells, matrix swelling and degradation, and cell-matrix remodelling are additional dynamic processes that take place post-fabrication and are capable of modifying the physico-chemical state of a bioprinted tissue model. The ability to map structural and mechanical characteristics of 3D tissue models non-destructively during and post-fabrication will present a significant asset to bioprinting technology, enabling bio-ink refinement and better understanding of artificial living systems’ functioning.

Brillouin microscopy could be just the right technology to meet this need. Evolved from the subfield of Brillouin spectroscopy, and initially applied to mechanical characterisation of solid state and crystalline materials in 1970s and 1980s [14, 15], Brillouin microscopy is now entering the field of biology and bioengineering with the promise to revolutionise micromechanical imaging of biomaterials, cells, tissues and organs thanks to its non-contact and label-free nature [16-19]. In Brillouin microscopy, the viscoelastic parameters are retrieved by scanning the sample point-by-point with the beam of low intensity light [20], thus allowing to probe native environments across 3D volumes with no need for special sample preparation. As light-based technology, however, Brillouin microscopy is similarly affected by the absorption and scattering in turbid matter, thus limiting the penetration depth in tissues and cell-matrix constructs to a few hundred micrometres on average [21].

Here we demonstrate how drop-on-demand 3D bioprinting is complemented by the state-of-the-art imaging technology such as Brillouin microscopy to achieve 3D contact-free and label-free micromechanical characterisation of complex, multicomponent biomaterials post-fabrication. We show that Brillouin microscopy are able to resolve small differences in viscoelasticity between hydrogel components with varying polymer concentrations as well as geometrical and structural features of the complex printed hydrogel models. This demonstration paves the way for Brillouin microscopy to be utilised as a complementary real-time characterisation method to achieve unprecedented level of control in fabrication of *in vitro* biological models.

## 2. Materials and methods

### 2.1 Materials

Px01.00 (H1) and Px04.00 (H2) RASTRUM matrix, with a storage modulus of 0.8 and 1.5 kPa respectively, (Inventia Life Science), Phosphate Buffer Solution (PBS, ThermoFisher), 96-well microplates (Nunc, ThermoFisher), 70 v/v% ethanol (Sydney Solvents) and DI water (Baxter) were used as received.

### 2.2 Hydrogel 3D bioprinting

All 3D hydrogel structures were designed using RASTRUM software and printed using RASTRUM digital 3D bioprinter (Inventia Life Science). The difference in the hydrogel composition stands in its total solid content, with relative difference between the two hydrogels being approximately 4.6%. The printing process followed the provided protocol by the supplier. Briefly, frozen bio-inks and activators were allowed to warm to room temperature for 40 minutes. A sterile cartridge was then filled with 70 v/v% ethanol (6 mL), sterile DI water (20 mL), and the defrosted bio-inks and activators. An automated sterilisation sequence was executed, which then followed by the priming of both the bio-inks and activators. The printing of the 3D structures was conducted using the recommended printing pressures for each ink. All structures were printed in a flat plastic-bottom, 96-well microplate.After printing, 150 µL PBS was added into each well and the plate was stored in the fridge until analysis.

Four different structures were bioprinted in this study. The single hydrogel square structure measured 3 mm in length and width, and 5 layers thick, printed into a 96-well microplate using either Px01.00 or Px04.00 hydrogel. The multi-hydrogel tri-lane structure with lane width of 800 µm and a single layer thickness utilized both Px01.00 and Px04.00 in one structure printed alternatingly. The multi-hydrogel, tri-layer structure also utilized both Px01.00 and Px04.00 in one structure, printed alternatingly. Each layer measured 3 mm in length and width, and 5 layers thick. Finally, the small plug dome structure has a diameter of 1.5 mm and 5 layers thick, formed using either Px01.00 or Px04.00.

### 2.2. Hydrogel Solid Content Measurement

Px01.00 hydrogel (200 µL) was formed manually in a tared petri-dish by transferring 100 µL of Px01.00 bio-ink, followed by 100 µL Px01.00 activator pipetted on top of the bio-ink. The petri dish was then covered with aluminium foil and transferred into a fume hood for 72 h to evaporate all the liquid. After evaporation, the mass of the remaining solid was measured and the solid content of the hydrogel was calculated according to: 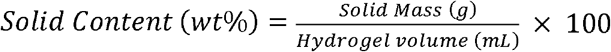. The process was repeated with the bio-ink and activator of Px04.00 to determine the solid content of Px04.00 hydrogel.

### 2.3 Brillouin microscopy

Brillouin microscopy was achieved through a combination of a confocal microscope with a specialised Brillouin spectrometer based on 6-pass tandem scanning Fabry-Perot interferometer (TFP1, JRS Instruments) as shown in Fig. 1a. A continuous frequency laser (660 nm, 120 mW, Torus, Laser Quantum) was used for sample illumination. The laser beam was coupled to a confocal microscope (CM1, JRS Instruments) and focused into the sample by a long working distance (WD) microscope objective (Mitutoyo, NA=0.42, WD=20 mm). After interacting with the sample (Fig. 1b), the light scattered in backwards direction was collected by the same objective lens and redirected to the 6-pass scanning Fabry-Perot interferometer for analysis. A spectrum of a hydrogel sample with Px01.00 hydrogel is shown in Fig. 1c: two side-bands (Stokes signal at -Ω and Anti-Stokes signal at Ω) are produced as the result of opto-acoustic interaction within the gel [17]. These bands are positioned symmetrically around the laser frequency which is manually set to 0 GHz. The exact position of Stokes and Anti-Stokes signals determines the Brillouin frequency shift (BFS, Ω), which is one of the two measurable outputs of Brillouin microscopy. For Px01.00 hydrogel samples, the BFS is found to be Ω_G_=6.05±0.005 GHz. The average and standard deviation for the Brillouin frequency shift are obtained based on averaging 5 measurement repeats for each hydrogel sample.

**Fig. 1.**
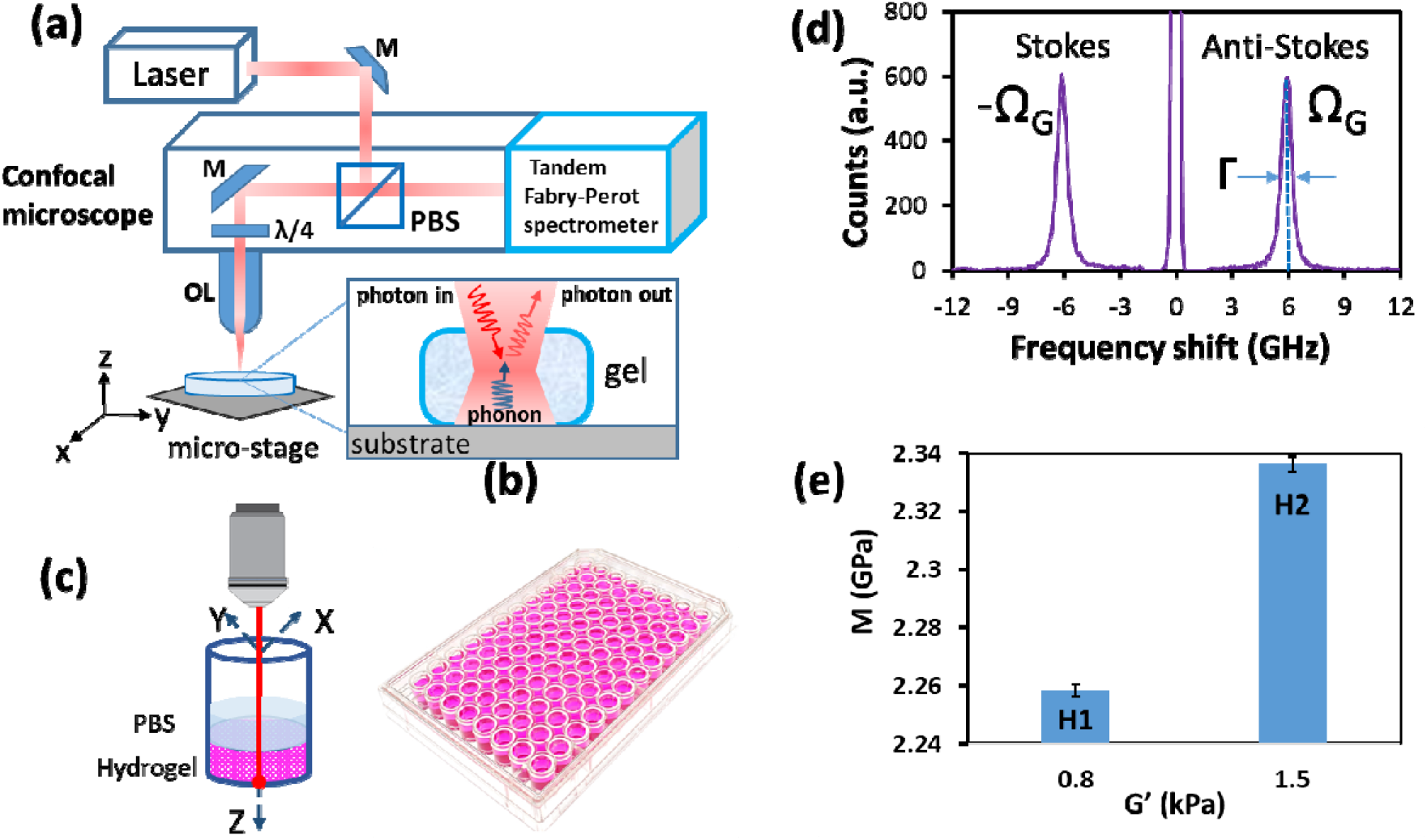
(a) Schematic of Brillouin microscopy setup. The 660 nm continuous wave laser is coupled with a confocal microscope and a tandem Fabry-Perot spectrometer to achieve GHz resolution imaging of gel micromechanical properties. M - mirror, PBS – polarisation beam splitter, OL - objective lens. (b) Schematic of photon-phonon interaction in a gel sample - as the result of such interaction, energy is either transferred from light to sound wave giving rise to Stokes signal (-Ω), or from sound to light giving rise to Anti-Stokes signal (Ω). (c) Schematic of a 96-well plate and a single well with hydrogel and PBS. (d) The measured spectrum of bioprinted hydrogel shows Brillouin frequency shift of Ω_G_=6.05±0.005 GHz. (e) Correlation between longitudinal (M) and shear (G’) moduli for samples with Px01.00 (H1) and Px04.00 (H2) hydrogel measured by Brillouin microscopy and rheology, respectively.

The linewidth of Brillouin Stokes and anti-Stokes signals (Γ) is the second measurable output of Brillouin microscopy. Both BFS and linewidth are directly related to the microscopic viscoelastic properties of the material. BFS can be expressed in terms of the longitudinal storage modulus (M), the refractive index of the material (n), the material density (ρ), the wavelength of light (λ) and the scattering angle (Θ) as 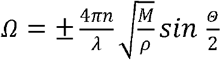 [17]. Thus, Ω is not only the function of the longitudinal modulus and stiffness but also depends on the refractive index and material density. It has been confirmed, however, that the ratio 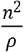 stays approximately constant in a wide range of biomaterials [22]. The Brillouin linewidth, on the other hand, is proportional to the material viscosity η given by 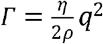 where 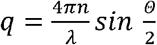 is the acoustic wave vector [16, 17]. For a backscattering geometry (*Θ* = 180°), the wave vector is 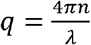 and the Brillouin frequency shift is 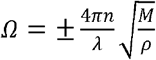.

The spectral resolution of the Brillouin microscopy system is mostly determined by the mirror spacing (L=4 mm) of the two tandem Fabry-Perot interferometers and for this set of experiments is 350 MHz (Fig. S1, see Supplementary information for details). Spatial resolution is determined by the objective lens, possible aberrations within the confocal microscope and properties of the spectrometer (the size of the input and output apertures). Taking this into account, the lateral resolution (resolution in XY plane) for our system is estimated to be approximately 2 μm. The axial resolution is measured by detecting Brillouin signal strength while scanning with the focused light beam across the boundary between two materials of distinctly different mechanical properties (e.g. plastic and water). The result of this test is presented in Fig. S2 in Supplementary materials and gives an estimate of the axial resolution to be approximately 120 μm. Two reasons contribute to this rather low axial resolution: 1) the type of the objective chosen for the measurement (the long distance objective with reasonably low numerical aperture NA=0.42) and 2) a large input aperture of the spectrometer (d=300 μm). The latter is of particular importance, since this degrades confocal nature of the microscopy system, meaning that the light scattered off focal plane can still reach the spectrometer and be added to the measured signal. To bring the system to the true confocal mode of operation, the aperture should be reduced down to approximately 25 μm, leading to severe reduction in the collected backscattered signal amplitude that in reality is not practical.

### 2.4 Data collection and processing

Brillouin microscopy data was collected using commercial (Ghost, JRS Instruments) and in-house built software to perform point measurements as well as l, 2 and 3D scans of BFS and linewidth. The same signal processing algorithm was applied to all data and consisted of two steps: 1) fitting of the measured data using Lorentzian line shape model to find the exact position of Stokes and Anti-Stokes peaks, and 2) compensating possible drift of the spectrum’s origin (laser frequency position) by averaging the data between Stokes and anti-Stokes peaks. Each point measurement was taken over acquisition time of 1-20 seconds. 20s long acquisition times were used to optimise signal-to-noise-ratio and improve the fitting precision. Linear, 2D and 3D scans were performed by moving the sample on the 3D microscopy stage along one, two or 3 axis of the stage, while keeping the optical system and the objective lens stationary.

### 2.5 Statistical analysis

The data are presented as mean ± standard deviation (SD) across n=5 samples of each type. One-way Anova (analysis of variance) with Tukey’s posthoc test for pairwise comparison was performed to obtain statistical difference between samples. Statistical significance is designated with \**p < 0.05*.

## 3. Results

### 3.1 Polymer concentration and time-dependent changes in hydrogel viscoelasticity

Both hydrogels, Px01.00 (H1) and Px04.00 (H2) were used in this study. Prior to conducting Brillouin microscopy studies, the solid content of the hydrogels were determined to be 13.8 and 18.4 wt% for Px01.00 and Px04.00 respectively. The Brillouin frequency shift of Px01.00 and Px04.00 hydrogels were measured at random locations within the hydrogel samples immersed in a phosphate buffered solution (PBS) over time. An increase of 1-1.5% in BFS shift was observed over the 14 days incubation period (Fig. 2). We hypothesised that the observed trend was due to a combination of changes in pH level, hydrogel swelling, molecular exchange between the gel and the PBS [23]. Further investigation into exact chemical changes is required to understand the time-dependent evolution of the hydrogel structure and composition, which is beyond the scope of the current work. Importantly, the BFS of distilled water did not show appreciable change across the 14-day measurement (p>0.05) confirming the stability of the measurement system over the incubation time.

**Fig. 2.**
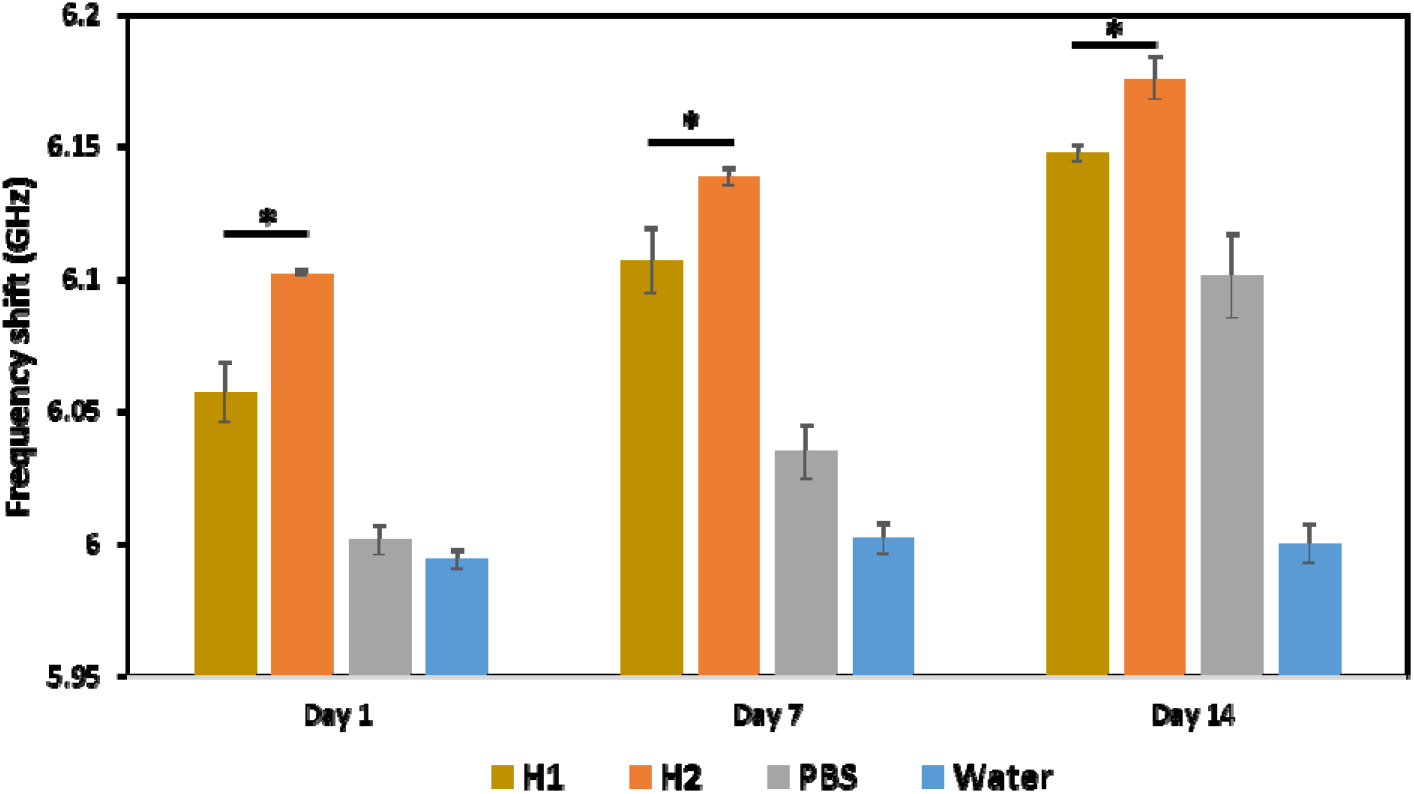
Brillouin frequency shift of Px01.00 (H1) hydrogel, Px04.00 (H2) hydrogel, PBS, and water measured *in situ* after incubation in PBS for 1, 7 and 14 days. Results are means ± SD.

More importantly, the BFS value for the Px04.00 hydrogel was higher than that for the Px01.00 hydrogel by approximately 50 MHz. This was attributed to the higher stiffness of the Px04.00 hydrogel due to an increase in crosslinking density and solid content [24]. The BFS differences of the two hydrogel compositions were consistent at each time point throughout the 14-day trial with stiffer gel showing the highest shift of Ω_G_ =6.1-6.17 GHz on average. Statistical analysis has confirmed the significance of this result (p<0.05). This demonstrates the sensitivity of the Brillouin microscopy technique, showing its capability to detect a small (4.6 wt%) difference in gel solid content. Importantly, we were not able to record significant difference in the Brillouin linewidth between Px01.00 and Px04.00 gels (see Fig. S3 in Supplementary material), meaning that possible difference in viscosity between the two gels is below the measurement capabilities.

To exclude ambiguity related to the continuing drift of the BFS in hydrogels and PBS over time, for the rest of experimental work reported here we used a relative BFS definition that takes into account the difference between the BFS of hydrogels and surrounding PBS, i.e.

### 3.2 Horizontal lanes patterns

Mimicking complex *in vivo* physiological cell environments and behaviours for *in vitro* applications requires the high-throughput production of multi-material 3D cell culture models. Such models comprise precise arrangement of hydrogels with different physical and biochemical properties to accommodate the desired response and growth of multiple different cell types within a high-content microliter plate. To test the capability of Brillouin microscopy on multi-materials hydrogels, we designed and RASTRUM printed multi-material tri-lane structures and multi-material tri-layer structures in a 96-well microplate.

The tri-lane structure comprised alternating hydrogel lanes with a lane width of 800 µm as shown in Fig. 3a. The results of scanning BFS across the lane pattern with the step size of 50 μm are presented in Fig. 3b. We found that the two hydrogel formulations with different stiffness and solid content were clearly distinguishable from each other and the PBS. The difference between BFS values of the hydrogels and PBS was found to be over 80 MHz, at least 8-fold larger than the measurement uncertainty of 9 MHz, estimated as the standard deviation across the data set for this measurement. The difference between the values of relative BFS between the two hydrogel compositions Px01.00 and Px04.00 was approximately 40 ±9 MHz.

**Fig. 3.**
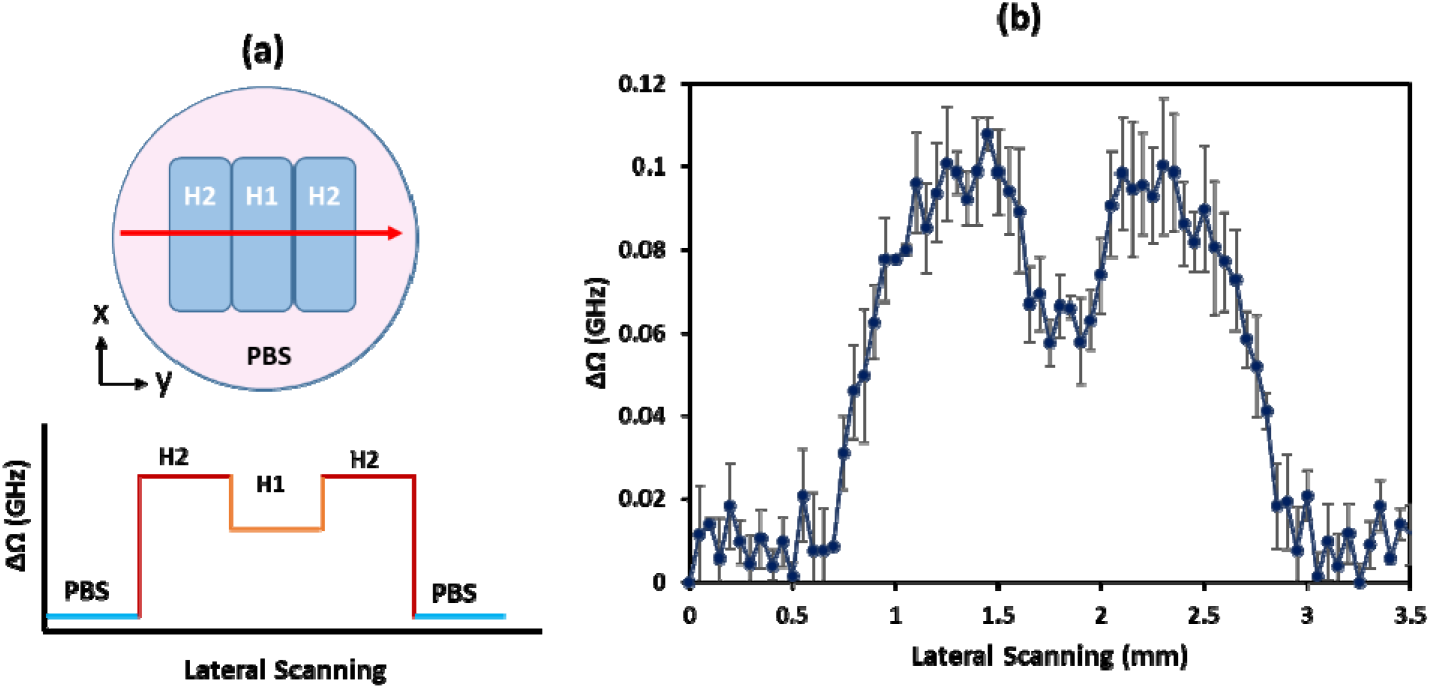
(a) The schematic of the multi-hydrogel tri-lane structure printed using Px01.00 (H1) and Px04.00 (H2) hydrogels and the expectation of a corresponding BFS measurement. (b) Lateral scanning of relative BFS for 800 μm wide hydrogel lane structures. Data stand for mean ± SD.

### 3.3. Vertical layer patterns

Next we printed a tri-layer structure of alternating hydrogel formulation to characterise the technique’s capability in micromechanical mapping along the z-direction. Scanning of the printed samples was conducted from the bottom of the well upwards in the direction orthogonal to the well surface, as schematically depicted in Fig. 4a. The estimated stack layer thickness was approximately 400 μm, thus we used a scanning step size of 50 μm to obtain a reasonable number of data points within each hydrogel layer. The hydrogel layers were clearly distinguishable from each other with the difference between relative BFS values of the hydrogel and the PBS of around 100 MHz (Fig. 4b). Similar to previous results discussed in Section 3.2, the BFS contrast between the two hydrogel formulations was found to be 50 ± 9 MHz.

**Fig. 4.**
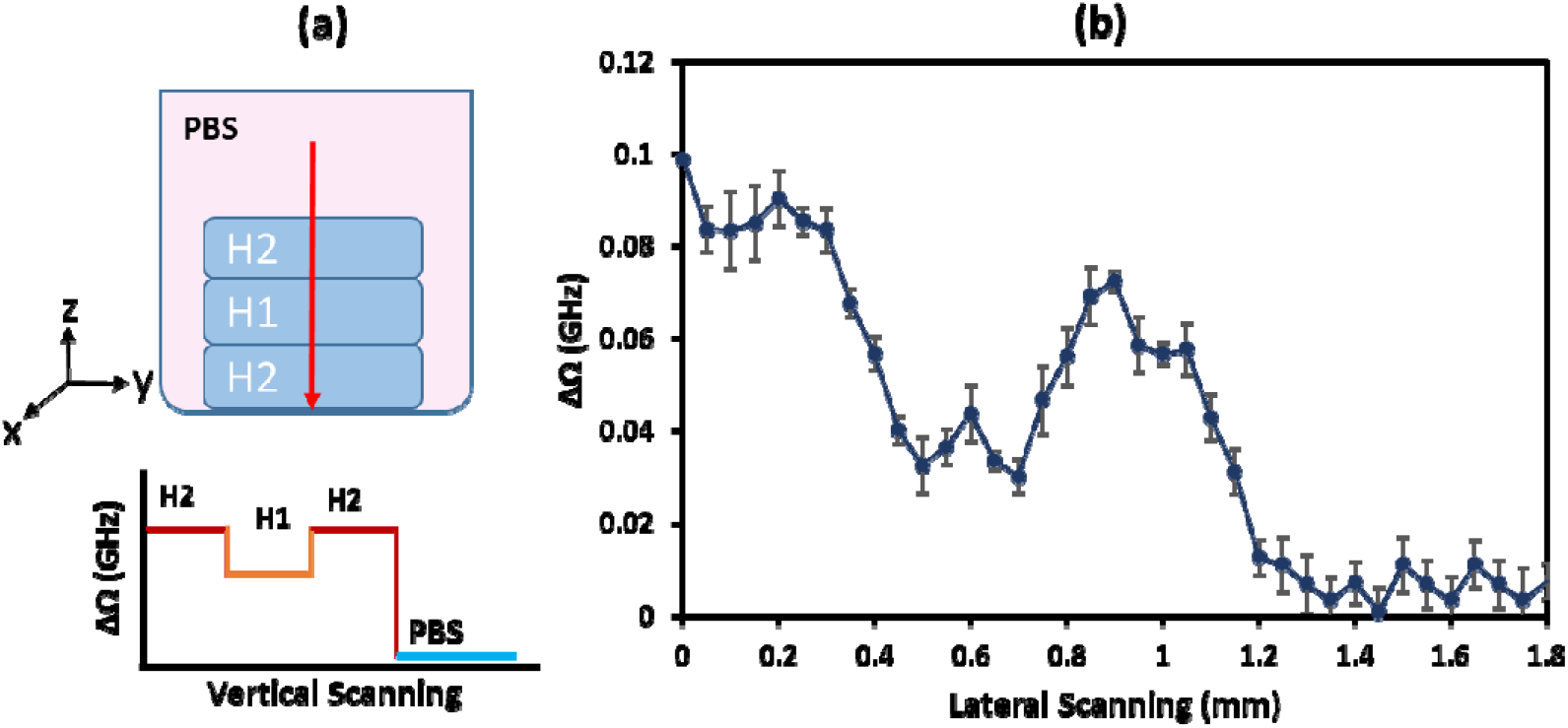
(a) Schematic of bioprinted multi-hydrogel vertical tri-layer structures and anticipated result of the Brillouin frequency shift measurement. (b) Measurement of the relative BFS across the stack with Px04.00 (H2) - Px01.00 (H1) - Px04.00 (H2) hydrogel layers. Data stand for mean ± SD.

The results obtained show that the resolution of the Brillouin microscopy technique is sufficient to spatially resolve multi-material 3D structure patterns made of multiple hydrogels with varying stiffness. Additionally, the size of each 3D structure post-fabrication and incubation in PBS can also be estimated from the BFS signals (subject to limitation in axial resolution of the measurement) and compared to the original design. For example, the height of each layer in multi-hydrogel vertical tri-layer structure shown in Fig. 4a and 4b was estimated to be approximately 300 μm. The knowledge of structural architecture and dimensions of multicomponent hydrogel models obtained from Brillouin microscopy measurements are of particular importance because of the hydrogel transparency in PBS that makes optical imaging of hydrogel structures unfeasible.

### 3.4 3D Brillouin imaging of the small plug structure

A small plug, dome-shaped 3D structure is a commonly used model for high-throughput drug screening application [25]. To mimic this, we RASTRUM printed a dome-shaped, 3D small plug cell model with a diameter of 1.5 mm in a 96-well plate, using either the Px01.00 or Px04.00 (Fig. 5). 3D mapping of the BFS of these structures was achieved by scanning the 96-well plate in X, Y and Z direction on the microscopy stage, while keeping the optical system fixed. For demonstration purposes, we present 2D Brillouin scans of the small plug structures at a number of positions along vertical Z-axis, 200 µm apart from each other as shown in Fig. 5a for Px01.00 and Fig. 5b for Px04.00 hydrogels. The colour scale in Fig. 5 uses blue-green coding with blue referring to the lowest relative BFS within the measurement and green for the highest. It is clearly noticeable that the size of the small plug structure was not uniform across Z direction, with the top of the structure being narrower compared to its bottom, reflecting the intended dome shape. The same can be said about BFS, with the bottom of the small plug structure showing higher BFS (brighter green) than the top. The average relative BFS and the standard deviation from the average for each layer are summarised in Table 1. On the z-axis, a decreasing relative BFS from the bottom to the top layer was observed, indicating a softer interface region than the bulk region of the hydrogel [26]. Correlating the BFS to storage modulus value using the calibration curve in Fig S4, we observed that the bulk region (G’ = 1.12 kPa) was around 1.3x stiffer than the interface (G’ = 0.87 kPa). Similar analysis on the BFS results across all layers of both Px01.00 and Px04.00 revealed on average a percentage BFS variance of ∼30% and ∼16%, respectively. The variance within these hydrogels was narrower than a swollen bioprinted Gel-Ma hydrogel having ∼50% variance [26] and a manually prepared Matrigel with ∼50% variance [27].

**Fig. 5.**
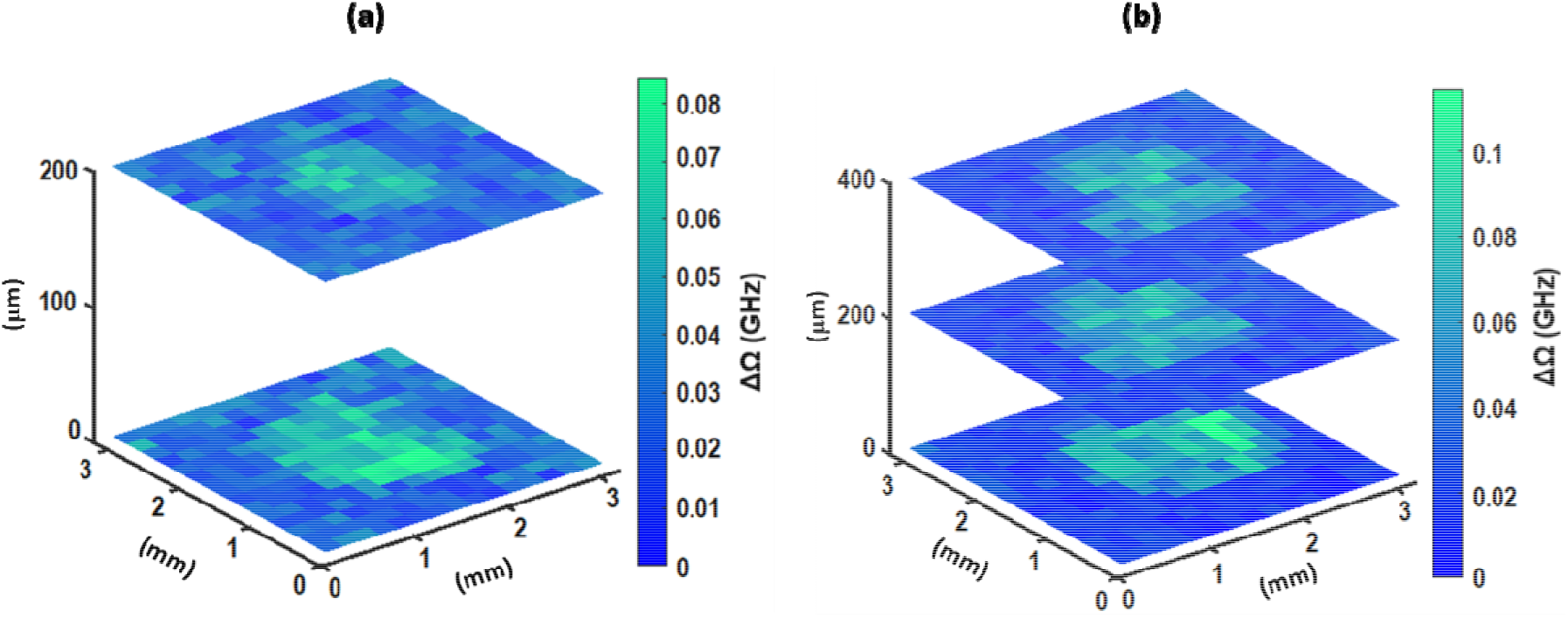
3D scanning of the relative BFS for (a) Px01.00 (H1) and (b) Px04.00 (H2) hydrogel small plug structure.

**Table 1.** Average relative BFS and standard deviation for Px01.00 (H1) and Px04.00 (H2) hydrogel small plug structures calculated based on images shown in Figs. 5a and b. PBS area outside the plug was excluded from the calculation.

|  | H1 | H2 |
| --- | --- | --- |
| Bottom layer | 0.035±0.012 GHz | 0.077±0.013 GHz |
| Middle layer | 0.026±0.008 GHz | 0.063±0.009 GHz |
| Top layer | - | 0.060±0.011 GHz |

As expected, Px04.00 small plug structure had a higher maximum BFS than that of the Px01.00. The observed heterogeneity in the elastic properties of the small plug structure can be explained if we consider the bioprinting technology and the mechanical variability that could be introduced throughout the crosslinking process. Taking into account that the crosslinking process depends on local stress distribution that would be quite different for bottom and top hydrogel layers, this is perhaps not surprising that the relative BFS is not constant across the 3D construct, meaning that the bottom of the small plug structure has higher longitudinal modulus and hence higher stiffness than the top of the structure.

Another important factor to consider is hydrogel swelling and the variation in the gel local hydration due to swelling. Since the small plug structure is attached to the bottom of the plastic well, its lower layers experience less swelling compared to its top layers that can expand freely. Thus, it is likely that the top of the structure has higher local hydration per unit volume, which would result in lower relative BFS.

In addition, the plug area is getting smaller by moving toward the top layer, which arises from the structure’s fabrication design. Brillouin microscopy, however, successfully reveals the spatial asymmetry and elastic heterogeneity of these visually transparent hydrogel constructs. It is worth mentioning that BM provides the same insight into the local micromechanical properties of hydrogel constructs as AFM microindentation but the measurement is much simpler, faster and truly compatible with imaging of 3D structures (versus AFM measurement of the 2D surface).

Overall, detected variability in micromechanical properties over 3D bioprinted hydrogel constructs suggests the important role of post-fabrication testing in order to precisely determine the structural imperfections and micromechanical distribution across bio-fabricated structures and be able to account for these imperfections in the interpretation of biological experiments.

## 4. Discussion

The presented results in this work reveal that BM can be successfully utilized to investigate the mechanical properties of hydrogel constructs through point measurement, line scanning and 3D mapping. The technique is sensitive to resolve differences in mechanical properties between two different hydrogels with small variations in polymer fraction and provides invaluable insight into hydrogel spatial distribution. Most importantly, the 3D mapping of transparent hydrogel constructs post-fabrication and media incubation is possible using this technique. This provides much needed insight into spatial uniformity of the hydrogel, gelation and swelling behaviour of the hydrogel models. This knowledge can be used to optimize printing protocols and improve the structure’s shape and mechanical properties.

As mentioned in Section 2.3, the confocal imaging volume is an oblate ellipsoid, which is elongated in Z-direction, and the 3D resolution of the microscope is 2 μm (X-direction) by 2 μm (Y-direction) by 120 μm (Z-direction). The axial resolution is extracted from the measurement of the axial spread function and discussed in more detail in Fig. S2 in the Supplementary information. This resolution is enough for the characterization of the 3D bioprinted scaffolds where the dimensions exceed a few hundred micrometres that is as presented in this work. Taking into account imaging resolution, the observation of BFS behaviour on the boundary between two different hydrogel materials or hydrogel and PBS presented in Figs. 3b and 4b requires additional clarification. Both scans across lateral and vertical directions show rather gradual boundaries for hydrogel structures, unlike the expected step-function behaviour illustrated schematically in Figs. 3a and 4a. Both scans were performed using 50 μm step size that is sufficiently larger than lateral imaging resolution (2 μm) and comparable to axial resolution (120 μm). Yet, the boundaries between both vertical and horizontal lanes are gradual indicating that another factor might be responsible for this effect rather than the spatial resolution of the system. We believe that mixing of the droplets printed at the interfaces and thus alterations to the crosslinking process and density could all be the possible explanations. This observation, however, indicates the importance of testing the hydrogel structures post-fabrication in order to compare the resulting architecture with the planned one, which would be especially important if the local mechanical properties of the hydrogel matrix determine cellular behaviour and, in some instances, even cell’ fate.

Overall, we have shown that Brillouin microscopy can be used to map micromechanical properties in hydrogel constructs that can be especially useful for scenarios of complex and inhomogeneous *in vitro* models. Another important application of Brillouin microscopy is in tracking dynamic changes in living systems post-fabrication. 4D bioprinting is an emerging field of research in which an additional dimension of time is added, suggesting temporal changes to chemistry, structure or physical properties of the model [28]. Swelling of the hydrogel constructs over time that is a desired effect and is programmed into the design to achieve time-dependent mechanical and chemical composition of the model is one of the possible 4D biomaterial examples. No matter the mechanism, Brillouin microscopy could be the key technology to confirm the desired temporal system behaviour or to provide the necessary feedback for the design adjustments.

In addition, Brillouin microscopy can be combined with Raman microscopy to add an extra channel of information and enable chemical mapping as well as mechanical. Non-contact and label-free nature of both Brillouin and Raman microscopy suggests that these technologies can be translated to *in vivo* imaging scenarios to harness their full potential [29-31]. Recent developments have also shown the possibility of fibre integration of Brillouin systems [32]. Thus in the future, it would be possible to integrate Brillouin fibre probe within a bioprinter nozzle to achieve simultaneous fabrication and characterisation of bioprinted scaffolds and cell models.

## 5. Conclusions

In conclusion, we have demonstrated that Brillouin microscopy can be utilised to probe the micromechanical properties of 3D printed hydrogel constructs. This technique provides non-contact and non-destructive approach for characterizing hydrogels with different stiffness and composition (solid fraction), as well as, revealing spatial structural and micromechanical distribution of complex hydrogel models. In addition, temporal changes such as swelling and degradation of the hydrogel constructs can be assessed to track evolution of hydrogel models over time.

## Acknowledgment

This work was supported by the Australian Research Council Discovery Program DP190110973.

## Appendix A. Supplementary data

